# Maternal regulation of the vertebrate oocyte-to-egg transition

**DOI:** 10.1101/2023.06.22.546137

**Authors:** Ricardo Fuentes, Florence L. Marlow, Elliott W. Abrams, Hong Zhang, Manami Kobayashi, Tripti Gupta, Lee D. Kapp, Zachary DiNardo, Felipe Montecinos-Franjola, William Vought, Charles Vejnar, Antonio Giraldez, Mary C. Mullins

## Abstract

Maternally-loaded factors in the egg accumulate during oogenesis and are essential for the oocyte and egg to acquire developmental competence and ensure the production of viable embryos. The oocyte-to-egg transition consists of the regulation of multiple molecular processes both cytoplasmic and nuclear acting in the late oocyte during a process called oocyte maturation. However, the molecular nature and functional importance of factors acting at this stage remain poorly understood. Here, we present a collection of 5 recessive maternal-effect mutants identified in a zebrafish forward genetic screen that reveal unique molecular insights into the mechanisms controlling the vertebrate oviparous oocyte-to-egg transition. We identified critical cytoplasmic regulators of yolk globule formation and maturation that are essential for egg development and embryogenesis. Specifically, the maternal-effect genes, *over easy*, *poached*, *p33bjta*, and *black caviar* control yolk globule sizing and/or protein cleavage during oogenesis, likely through endolysosomal organization independent of nuclear oocyte maturation. Furthermore, we cloned one of the mutant genes, identifying a subunit of the Adaptor Protein complex 5, which regulates intracellular trafficking, and yolk vesicle formation. Together, these mutant genes represent novel genetic entry points to decipher the molecular mechanisms functioning in the oocyte-to-egg transition, fertility, and human disease. Additionally, our genetic screen provides valuable functional tools for exploring the evolutionary fates of maternal factors and their contribution to developmental strategies for reproductive success in metazoans.

**Author Summary:** The oocyte-to-egg transition consists of the coordinated regulation of multiple molecular processes acting in the late oocyte. This transcriptionally silent period requires the precisely timed function of maternally-supplied gene products during oogenesis. However, knowledge of their molecular nature and *in vivo* function remains incomplete. The mutants reported here provide access to maternal factors regulating the processes that prepare an oocyte for reproductive competence and embryogenesis. We have identified essential regulators of yolk granule formation and protein processing. Specifically, we found that the highly conserved maternal Ap5m1 protein regulates yolk granule maturation, which generate essential nutrients and immunity for growth and development in oviparous animals. The mutants presented here represent attractive genetic models to investigate the molecular and cell biological mechanisms that control the oocyte-to-egg transition, as well as reveal a collection of genetic factors indispensable for reproduction and survival. Importantly, knowledge of their genetic underpinnings and biological importance in reproduction will also pave the way to investigate the evolution of maternal genes during vertebrate development.

## Introduction

Animal embryonic development depends on factors endowed to the oocyte during oogenesis. These factors make the oocyte competent to mature into an egg that will activate and subsequently undergo embryonic development by a process called the oocyte-to-embryo transition. This gradual and tightly regulated developmental program requires the function of gene products, called maternal factors, that are loaded into the egg during oogenesis (reviewed in Abrams & Mullins, 2009; Horner & Wolfner, 2008; Lindeman & Pelegri, 2010; Robertson & Lin, 2013; Schultz, Stein, & Svoboda, 2018). Oocyte maturation and egg activation are fundamental to the oocyte-to-embryo transition, but how these events are molecularly regulated is not well understood. In fact, the molecular identity and function of most of the maternal factors orchestrating the embryonic program in vertebrates remain unknown, highlighting the need for developing methods to identify and study them.

The oocyte undergoes growth and maturation during early and late stages of development, respectively, to transition into an egg. This involves changes in the function of specific genes, the regulation of meiosis, and cellular signaling pathways that are necessary for successful fertilization and early embryonic development (He, Zhang, Yang, & Wang, 2021; Hsu, Pan, Wang, Tong, & Chung, 2018; Lessman, 2009; Wallace & Selman, 1990)}. Research on oocyte maturation is a rapidly advancing field and has implications for understanding fertility, reproductive disorders, and the development of assisted reproductive technologies (ART)(Dvoran, Nemcova, & Kalous, 2022; Innocenti et al., 2022).

The oocyte-to-egg transition encompasses large-scale cytoplasmic changes, preparing an oocyte capable of being fertilized by sperm (Selman et al., 1993). Early cytoplasmic reorganization includes animal-vegetal axis establishment, the formation of secretory vesicles or cortical granules (CGs), and vitellogenesis, where yolk proteins are synthesized in the liver, transported to the ovaries, and taken up by the developing oocytes (Eppig, 1996; Fuentes et al., 2020; Rojas et al., 2021; Romano, Rosanova, Anteo, & Limatola, 2004; Stitzel & Seydoux, 2007). In many oviparous organisms, including zebrafish, the oocyte accumulates maternal Vitellogenin (Vtg) protein via receptor-mediated endocytosis (Wallace & Selman, 1990). Maternal Vtg, a conserved phospholipoglycoprotein, is the primary yolk protein and a vital nutrient for embryonic development in many egg-laying species, including platypus, but not placental mammals (Brawand, Wahli, & Kaessmann, 2008). Once Vtg is internalized, it is partially processed in endosomes and then stored as yolk globules (YGs) during oogenesis (reviewed in (Fagotto, 1995). YGs are modified lysosomes and contain hydrolases that process the yolk proteins; however, unlike typical lysosomes, YGs do not immediately process their contents (Fagotto, 1995; Wall & Meleka, 1985). As important lysosomal vesicles, YGs and the mechanism behind their formation and function could have clinical implications, since the etiology of most vesicle trafficking- and lysosomal-related defects in human disease remain unresolved (reviewed in (Bonam, Wang, & Muller, 2019; Huizing, Helip-Wooley, Westbroek, Gunay-Aygun, & Gahl, 2008; Platt, Boland, & van der Spoel, 2012).

During late stages of oogenesis, nuclear maturation and cytoplasmic changes (cytoplasmic maturation) together produce a viable egg, including a shift in the metabolic and optical properties of YGs (Dosch et al., 2004; Eppig, 1996). In addition, a yolk-free and well-defined cytoplasmic domain accumulates at the animal pole, forming the preblastodisc, highlighting the relevance of the spatiotemporal and asymmetrical positioning of maternal fate-determining molecules and organelles within the oocyte (Fernández et al., 2006; Fuentes et al., 2018; Lessman, 2009; Selman et al., 1993). During oocyte maturation in vertebrates, the oocyte arrests in meiosis II until ovulation of the egg and its subsequent activation (reviewed in (Holt, Lane, & Jones, 2013; Lessman, 2009; Marlow, 2010; Voronina & Wessel, 2003). How these different aspects of cytoplasmic maturation are controlled in space and time is still largely unknown.

Here, and in a companion study, we report a forward genetic adult screen to identify novel maternal genes functioning in the oocyte-to-embryo transition in the zebrafish. We identified 5 maternal-effect mutants falling into a class of phenotype with defects in oogenesis, specifically in aspects of YG formation and maturation. This collection of vertebrate mutants represents a valuable resource for understanding the molecular regulation of the oocyte-to-egg transition, providing genetic entry points and models of infertility or poor reproductive success.

## RESULTS

### Identification of zebrafish oocyte-to-embryo transition mutants

To isolate mutations in genes regulating the oocyte-to-embryo transition in zebrafish, we induced mutations with the chemical mutagen N-ethyl-N-nitrosourea (ENU) and conducted a four-generation adult maternal-effect screen for egg and embryo phenotypes (Dosch et al., 2004; Wagner, Dosch, Mintzer, Wiemelt, & Mullins, 2004). We examined 716 mutagenized genomes, and identified 9 *bone fide* (recessive) oocyte-to-embryo transition maternal-effect mutants (Table 1) that failed to produce eggs competent to undergo early embryonic development. We identified an additional 9 maternal-effect mutants disrupting the cleavage stage of embryogenesis in zebrafish (Abrams et al., 2020). Five of the oocyte-to-embryo transition mutants displayed phenotypes indicating a primary defect during oogenesis (this work), including one new mutant allele of the *over easy* (*ovy*) gene (Dosch et al., 2004). Four additional mutant genes were identified that disrupt egg activation (accompanying paper).

**Table 1.** Oocyte-to-Egg Transition Zebrafish Maternal-effect Genes.

| Mutant Class | Gene/allele | Molecular identity<sup>a</sup> | Chromosomal Locus<sup>b</sup> |
| --- | --- | --- | --- |
| Oocyte-to-egg transition | *p33bjta* | ND | ND |
|  | *over easy*<sup>p35aluc</sup> | *ap5m1* | Chr 17, z7170 |
|  | *over easy*<sup>p37ad</sup> | *ap5m1* | Chr 17, z7170 |
|  | *poached*<sup>p26ahubb</sup> | ND | Chr 17, z21194 |
|  | *black caviar*<sup>p25bdth</sup> | ND | Chr 5, z3804 |
|  | *black caviar*<sup>p26ahubg</sup> | ND | Chr 5, z3804 |

ND= Not determined
(a) *ap5m1*= adaptor related protein complex 5 subunit mu 1
(b) Chromosome (Chr), closely linked marker

### Oocyte-to-egg transition mutant phenotypes and gene mapping

The largest group of mutants identified in our screen exhibits defects in the oocyte-to-egg transition (Fig. 1A). In zebrafish, the clarity of the yolk is a cytoplasmic indicator of oocyte maturation. Prior to oocyte maturation in wild-type, the yolk is opaque and the bulk of the cytoplasm is intermingled between the YGs in the oocyte. During oocyte maturation, the yolk proteins are cleaved, generating the translucency of the egg (Fig. 1A) (Selman, Wallace, Sarka, & Qi, 1993). Three mutants reported here, *p33bjta, p26ahubb,* and *p35aluc,* produced eggs that were opaque and lacked a blastodisc, more closely resembling wild-type oocytes than eggs (Fig. 1A). The two other mutants of this class, *p26ahubg* and *p25bdth* (hereinafter called *black caviar^p25bdth^* (*blac^p25bdth^*), only produced degenerated eggs (Figs. 1B, S1A), indicating a more severe or distinct defect in the oocyte-to-egg transition.

**Figure 1.**
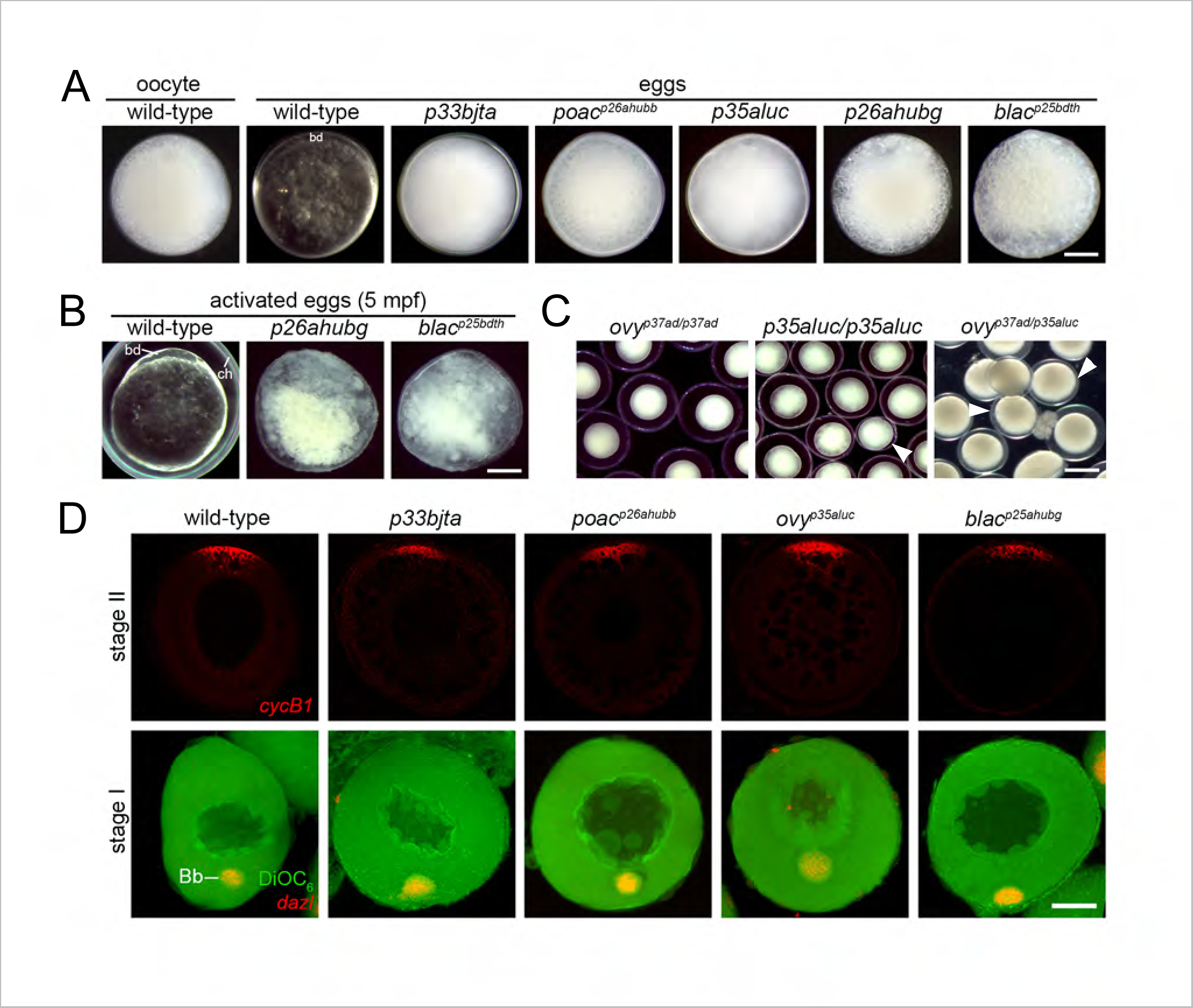
Zebrafish oocyte-to-egg transition mutants. **A.** Wild-type stage IV oocyte and egg in incident light. Eggs of *p33bjta* (1570 from 21 females)*, poac^p26ahubb^* (1620 from 35 females), and *p35aluc* (2088 from 25 females) resemble wild-type oocytes in their opacity, while eggs of *p26ahubg* (546 from 8 females) and *blac^p25bdth^* (1020 from 17 females) were degenerated. **B.** Images of 5 mpa wild-type (n=68), *p26ahubg* (n=56) and *blac^p25bdth^* (n=61) eggs, showing lysed mutant eggs. **C.** Non-complementation test showed that females carrying the *ovy^p37ad^* mutation in trans to the *p35aluc* mutation lay opaque mutant eggs (n=1382 from 9 females), thus confirming that *p35aluc* is an *ovy* allele. **D.** Confocal z-projections of isolated acid fixed oocytes analyzed by fluorescent *in situ* hybridization and also DiOC_6_ staining (green) in lower panels. In stage II oocytes, *cycB1* transcript localized to the animal pole in wild-type (n=32), *p33bjta* (n=22), *poac^p26ahub^* (n=47), *ovy^p35aluc^* (n=38), and *blac^p25bdth^*(n=28) oocytes. In stage I oocytes, *dazl* mRNA co-localized with DiOC6 to the vegetal Balbiani body (Bb) in wild-type (n=32), *p33bjta* (n=23), *poac^p26ahub^* (n=63), *ovy^p35aluc^* (n=31), and *blac^p25bdth^* (n=29) oocytes. Scale bar= 180 µm (A), 200 µm (B), 700 µm (C).

To determine if the oocyte-to-egg transition mutations represent alleles of the same or distinct genes, we determined the chromosomal positions of each mutation through bulk segregant analysis using simple sequence length polymorphic (SSLP) markers (Knapik et al., 1998). The *p35aluc and p26ahubb* mutations mapped to a similar chromosomal location as the *over easy (ovy^p37ad^)* mutation, which also causes an opaque egg phenotype (Dosch et al., 2004). To investigate if *p35aluc* or *p26ahubb* is allelic to *ovy^p37ad^,* we conducted complementation tests. We found that the eggs of *p35aluc/ovy^p37ad^* females were opaque, similar to that of either the *ovy^p37ad^* or *p35aluc* homozygous mutant females (Fig. 1C). Based on linkage of *p35aluc* and *ovy^p37ad^* mutations to the same interval on chromosome (Chr) 17 and the genetic non-complementation, we conclude that *p35aluc* is an allele of *ovy^p37ad^* (hereafter called *ovy^p35aluc^*). We performed similar complementation analysis for *p26ahubb* and *ovy^p35aluc^*, crossing two heterozygous *p26ahubb* females to two homozygous *ovy^p35aluc^*males. No females from the progeny displayed the opaque egg phenotype (0/36 and 0/38 females). Thus, unlike *ovy^p35aluc^* and *ovy^p37ad^, p26ahubb* complemented *ovy^p35aluc^*, indicating that the *p26ahubb* mutation disrupts a different gene, which we named *poached* (Table 1, hereafter called *poac^p26ahubb^*). Among the mutants identified in this screen, only *p35aluc* represented a new allele of a previously reported zebrafish maternal-effect gene.

Homozygous females of the *blac^p25bdth^* and *p26ahubg* mutations produced an identical phenotype, lysed eggs at egg laying (Figs. 1B; S1A). We mapped the *blac^p25bdth^*mutation to Chr 5 and then determined that the *p26ahubg* mutation was linked to the same SSLP marker (Table 1), suggesting that they are mutations in the same gene. We performed complementation analysis between these two mutations, crossing three *p26ahubg* heterozygous females to either *p25bdth* homozygous males (n=2) or a heterozygous male (n=1). We raised the fish to adults and tested females for the lysed egg phenotype. We found that these mutations failed to complement each other with mutant females from all 3 crosses in the expected ratios (10/20, 15/31, and 2/10 females for the three respective crosses). Thus, these two mutant alleles disrupt the *blac* gene.

We did not find linkage between the *p33bjta* mutation and the SSLP markers tested. However, we have propagated the mutation through many generations and our analysis indicates that the *p33bjta* mutation is not linked to the chromosomal interval of any of the other opaque egg mutations reported here or previously (Dosch et al., 2004), suggesting that it disrupts a distinct gene.

### Oocyte-to-egg transition mutants exhibit defects in egg activation

All mutants of this class produced eggs that failed to form the blastodisc at the animal pole and showed no overt signs of egg polarity (Figs. 1A-C). Therefore, we examined markers of the animal-vegetal axis to investigate whether failed blastodisc formation was due to a defect in oocyte polarity. We found that like in wild-type, in oocytes of *p33bjta*, *ovy^p35aluc^, poac^p26ahubb^* and *blac^p25bdth^,* c*yclinB1* (*cycB1*) and *deleted in azoospermia-like* (*dazl*) mRNAs localized to the animal pole of stage II and to the vegetal Balbiani body of stage I oocytes, respectively (Fig. 1D). Therefore, impaired oocyte polarity does not cause the lack of animal-ward segregation of cytoplasm that forms the blastodisc.

We also investigated whether mutant eggs completed meiosis following egg activation by examining second polar body formation (Dekens, Pelegri, Maischein, & Nusslein-Volhard, 2003). We found that meiosis resumed normally at 5 minutes post-egg activation (mpa) in the mutants, with sister chromatids evident in anaphase II (Fig. 2A). By 10 mpa, an actin associated second polar body was detected in wild-type and mutant eggs (Fig. 2B), indicating normal completion of meiosis II.

**Figure 2.**
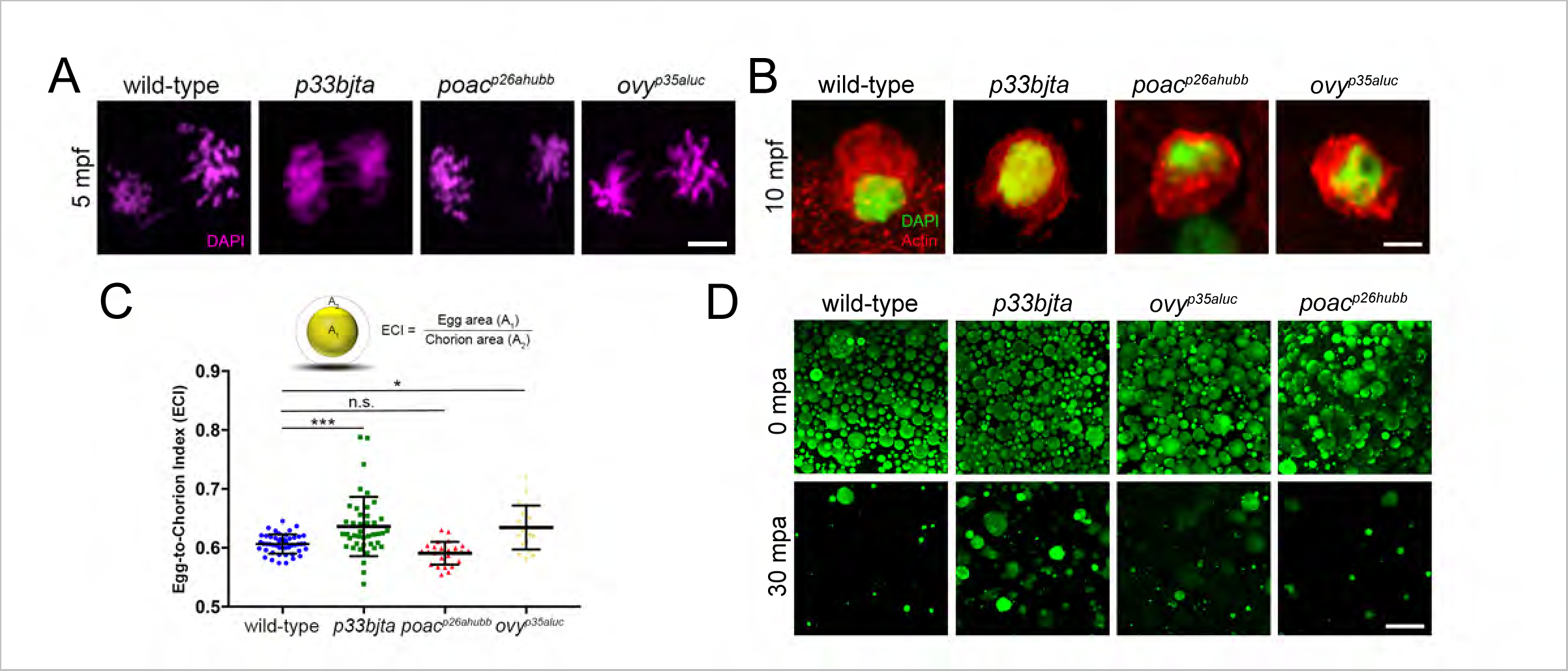
Meiosis II completion in dark egg mutants. **A.** Confocal micrographs of the animal pole show anaphase of meiosis II at 5 mpa and the extruded polar body at 10 mpa. DAPI-stained wild-type (n=24), *p33bjta* (n=14), *poac^p26ahubb^*(n=22) and *ovy^p35aluc^* (n=21) sister chromatids during meiotic anaphase II. **B.** Phalloidin-stained actin (red) and DAPI staining (green) of second polar bodies in wild-type (n=14), *p33bjta* (n=9), *poac^p26ahubb^* (n=12) and *ovy^p35aluc^* (n=10) activated eggs. **C.** Egg-to-chorion index (ECI) is the ratio of egg area (A_1_) to chorion area (A_2_) as a measure of chorion elevation. Scatter plots of the ECI measured in activated eggs at 30 mpa. The wild-type egg ECI value (ECI=0.61; SD=0.02; n=65) defines the numerical value for normal chorion elevation. The ECI for the eggs from *p33bjta* (ECI=0.64; SD=0.05; n=44, 2 females) and *ovy^p35aluc^*(ECI=0.63; SD=0.03; n=20, 2 females) mutant females are significantly different to wild-type, while the eggs of *poac^p26ahubb^* (ECI=0.59; SD=0.02; n=25, 2 females) mutant females show a mean value similar to wild-type. **D.** Confocal z-projections (5 μm depth) of acid fixed and MPA stained wild-type and mutant activated eggs. In wild-type (n=60), *p33bjta* (n=55), *ovy^p35aluc^* (n=25), and *poac^p26ahubb^* (n=27) eggs, numerous large and small CGs were distributed through the cortex at 0 mpa. At 30 min after egg activation, wild-type (n=21), *p33bjta* (n=28), *ovy^p35aluc^*(n=18), and *poac^p26ahubb^* (n=20) eggs displayed fewer CGs, indicating that they were released following activation. However, some CGs persisted in *p33bjta* (5/28) and *ovy^p35aluc^* (6/18) mutant eggs, which may evoke the reduced chorion expansion observed in mutant eggs (Figs. 1C, 2C). SD=standard deviation. mpa, minutes post-activation. Data are means ± SD. \*\*\**p*=0.0004; \**p*=0.0131; n.s., not significant. Scale bar= 4 µm (A, B), 40 µm (D).

To address whether other aspects of egg activation were altered in the mutants, we investigated CG exocytosis and chorion elevation. We first defined the egg-to-chorion index (ECI) as a readout of chorion elevation. To calculate ECI, activated egg and chorion areas were measured at 30 mpa (see Materials and Methods). Quantitative analysis of the ECI of *poac^p26ahubb^*eggs showed normal chorion elevation (*p*>0.05) (Fig. 2C). However, *p33bjta* and *ovy* eggs displayed a significant, variably expressive reduction in chorion elevation (Figs. 1C and 2C). We next examined CG exocytosis, which mediates the chorion expansion, and found that small and large CGs were present and released after activation of wild-type and mutant eggs (Fig. 2D). However, some large CGs persisted in the *p33bjta* and *ovy* eggs at 30 mpa (Fig. 2D), which may cause the variably expressive chorion elevation phenotype.

These data show that features of egg activation are perturbed in these mutants. While the capacity to undergo meiotic divisions and polar body extrusion is normal, defective cytoplasmic segregation and additionally chorion elevation in some mutants suggest that specific egg activation traits are regulated by bifurcating or distinct genetic networks.

### Oocyte cytoplasmic maturation of YGs

The zebrafish stage III oocyte is opaque due to the accumulation of membrane-enclosed crystalline yolk; however, at the end of oocyte maturation the yolk becomes non-crystalline and translucent (Lessman, 2009; Selman et al., 1993). This process is driven by the proteolytic processing of the major yolk proteins during oocyte maturation in stage IV oocytes, changing their refractive index from opaque to translucent. The persistent opaque yolk in these mutant eggs suggests that cleavage of the major yolk proteins during oocyte maturation is impaired. To investigate this directly, we prepared extracts from wild-type eggs and immature stage IV vitellogenic oocytes and compared the yolk protein profiles to those of the oocyte-to-egg transition class of mutant eggs. We found that the yolk protein composition of mutant eggs of *blac^p25bdth^*, *poac^p26ahubb^*, *p33bjta* and *ovy^p35aluc^*resembled wild-type oocytes rather than wild-type eggs (Fig. 3A), indicating that cleavage of the vitellogenic proteins was indeed defective during oocyte maturation in these mutants.

**Figure 3.**
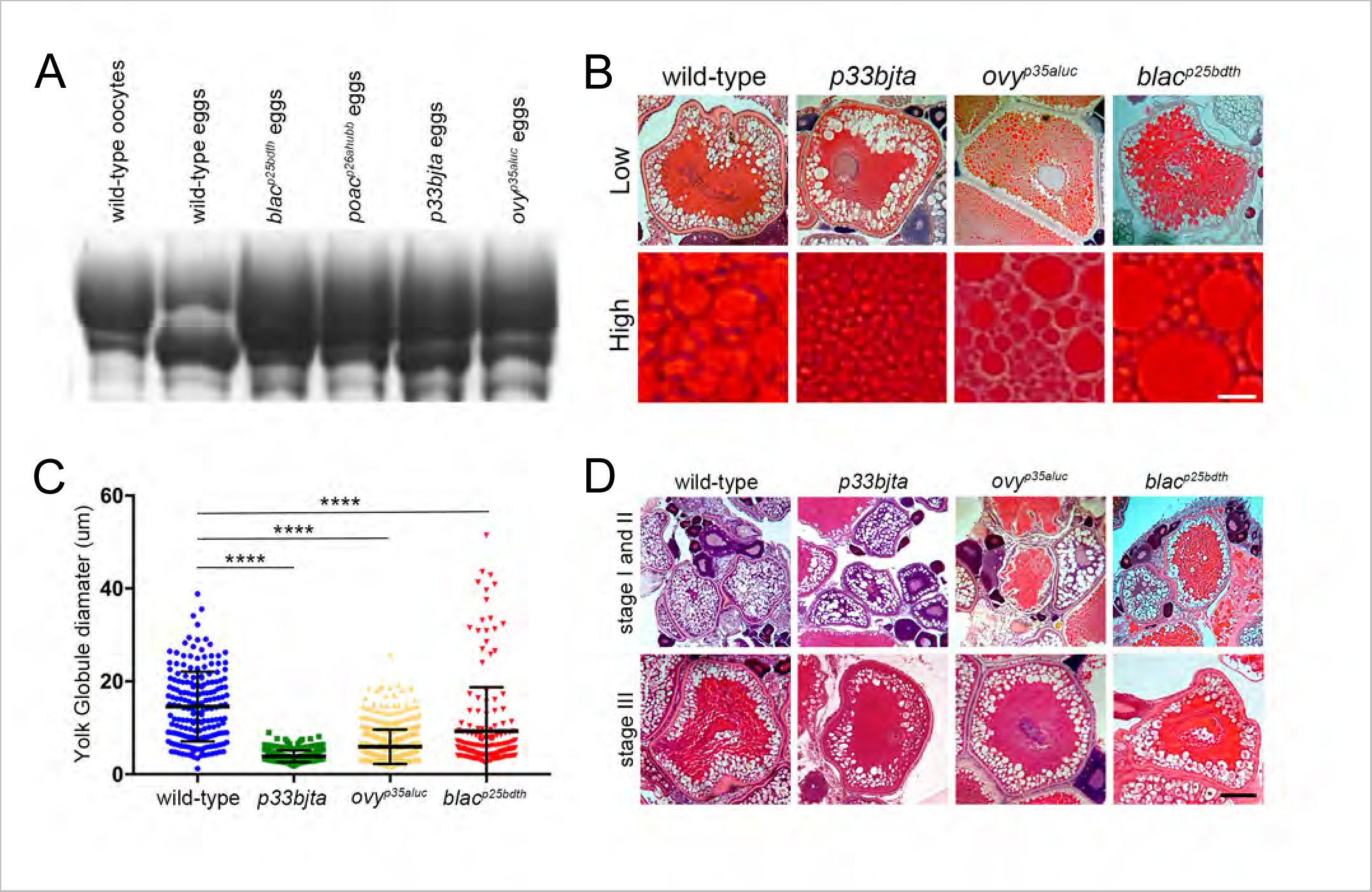
Proteinogenic and morphological features of the YGs in the oocyte-to-egg transition mutants. **A.** Coomassie stained gel of major yolk proteins in wild-type and mutant eggs showing that cleavage of yolk protein is impaired in the oocyte-to-embryo transition eggs. **B.** Low (top row) and high (bottom row) magnification histological sections of wild-type and mutant stage III oocytes stained with H&E revealing YG sizes. **C.** Scatter plots of YG diameter phenotype measured in a defined region of the oocyte. The mean diameter of the YGs in the mutants was significantly different to that of wild-type oocytes (WT=14.6 µm, *p33bjta*=3.87 µm, *p35aluc*=5.95 µm, *p25bdth*=9.25 µm). The number of YGs analyzed in this study was: wild-type (n=188 from 4 oocytes), *p33bjta* (n=320 from 2 oocytes), *ovy^p35aluc^* (n=503 from 2 oocytes), and *blac^p25bdth^* (n=177 from 2 oocytes). Data are means ± SD. \*\*\*\**p*<0.0001. **D.** Hematoxylin and Eosin stained sections of intact dissected ovaries (top row) and stage III oocytes (bottom row). Wild-type (n=12 ovaries), *p33bjta* (n=11 ovaries), *ovy^p35aluc^*(n=10 ovaries) and *blac^p25bdth^* (n=14 ovaries) were phenotypically comparable and all oogenesis stages can be found. Stage III oocytes were also comparable in size and morphology to wild-type oocytes. Scale bar= 16 µm (B), 95 µm (D).

To unravel the cellular mechanism underlying protein cleavage during YG maturation, we asked whether this process relies on YG morphology and integrity. We therefore examined the size of YGs in wild-type and mutant stage III oocytes in hematoxylin and eosin (H&E) stained histological sections (Fig. 3B). We found that the YGs of wild-type stage III oocytes had a mean diameter of 14.6 μm (Fig. 3C). In *p33bjta*, *ovy^p35aluc^* and *blac^p25bdth^* mutants, the YGs were on average 3.7-, 2.5-, and 1.6-fold smaller than in wild-type, respectively (Figs. 3B,C). The *p33bjta* phenotype was more severe than *ovy^p35aluc^*, displaying a much greater amount of small YGs and no large YGs (Fig. 3B,C). In *blac^p25bdth^* mutant oocytes numerous, smaller YGs were present, but some large YGs also formed, with a few larger than in wild-type (over 40 μm) (Figs. 3B,C). Thus, quantification of YG size confirmed a significant difference between wild-type and *p33bjta*, *ovy^p35aluc^*, and *blac^p25bdth^*mutants (Fig. 3C).

Histological sections of earlier stage I and II oocytes revealed no overt differences between mutants and wild-type (Fig. 3D). H&E staining indicated that previtellogenic oocytes were normal; cortical granules accumulated and later localized to the oocyte cortex in *ovy^p35aluc^*, *p33bjta*, and *blac^p25bdth^* mutant stage III oocytes (Fig. 3D).

These data indicate that YG size was severely compromised in *ovy^p35aluc^*and *p33bjta* mutants; thus, suggesting that the products of the *ovy* and *p33bjta* genes promote the formation of large YGs. Taken together, the anomalies observed in these mutants strongly suggest that *ovy*, *p33bjta*, and *blac* regulate the initial steps of YG formation and yolk protein processing, independent of other aspects of oocyte cytoplasmic and nuclear maturation.

### ovy encodes adaptor-related protein complex 5, mu 1 subunit (ap5m1)

To determine the molecular nature of the *over easy* gene, we performed RNA sequencing (RNA-seq) of *ovy^p37ad^* mutant eggs, an approach used previously to identify mutant genes in zebrafish (Hill et al., 2013; Langdon et al., 2016; Miller, Obholzer, Shah, Megason, & Moens, 2013; Reischauer et al., 2016). A search of the top ten downregulated genes in the mutant egg transcriptome identified only one gene in the chromosomally defined *ovy* interval on Chr 17, the *<u>a</u>daptor-related <u>p</u>rotein complex <u>5</u>, <u>m</u>u <u>1</u> subunit* (*ap5m1*) gene (log_2_ fold change= −7.0632) (Fig. 4A; see Materials and Methods). It encodes one of four subunits of the fifth Adaptor Protein complex (AP5) (Hirst et al., 2011), the mu or μ subunit. The AP5 heterotetramer complex localizes to a late endosomal/lysosomal compartment and functions in sorting late endosomes back to the Golgi in cultured mammalian cells (Hirst et al., 2011; Hirst et al., 2015; Hirst, Irving, & Borner, 2013; Hirst, Itzhak, Antrobus, Borner, & Robinson, 2018).

**Figure 4.**
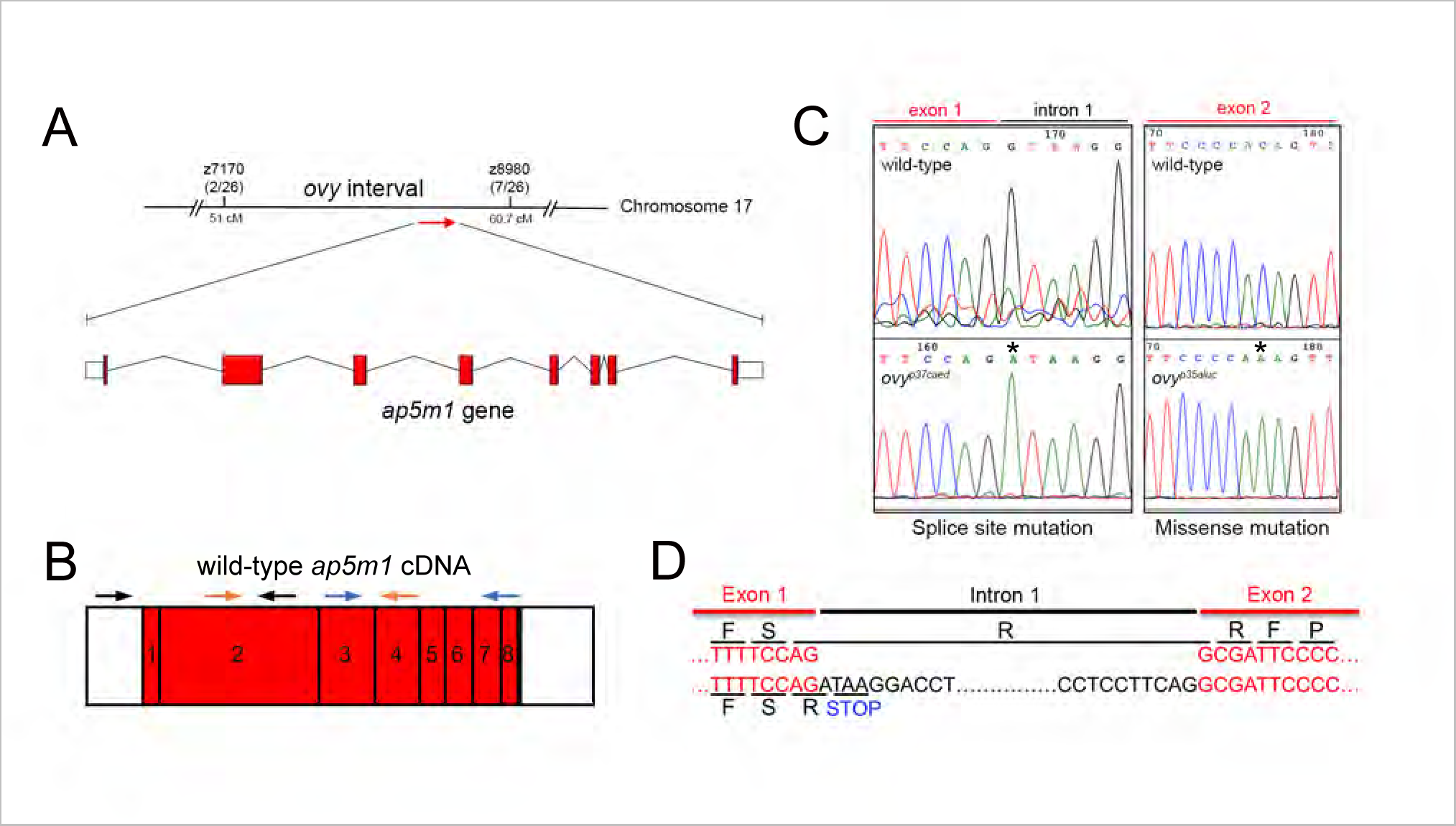
Molecular nature of the *ovy* gene. **A.** Genetic and physical map of the *ovy* locus and schematic representation of the zebrafish *ap5m1* gene, which consists of 8 exons and 7 introns. z7170 and z8980 are SSLP markers flanking the *ovy* mutation. In parenthesis the number of recombinants/analyzed meioses. Exons are shown as red boxes and introns as black lines. Sizes are not to scale. **B.** *ap5m1* cDNA amplification schematic. The relative positions of the different primers used are above the *ap5m1* gene. Three different color-coded pairs of primers are shown. Exons are represented as numbered red boxes. **C.** DNA sequencing analysis of the *ovy^p37ad^* and *ovy^p35aluc^*mutant alleles. Left: genomic DNA sequence of the wild-type and *ovy^p37ad^* allele indicates a single point mutation in the splice donor site of intron 1 of the *ap5m1* gene (black asterisk). Right: cDNA sequence of the wild-type and *p35aluc* allele indicates a single point mutation in exon 2 of the *ap5m1* gene (black asterisk). **D.** Insertion of intron 1 found in the *ovy^p37ad^* mutant allele. Predicted STOP codon encoded by the *ovy^p37ad^* mutant *ap5m1* transcript suggests that the mutant transcript would produce a structurally incomplete protein lacking the C-terminal portion of the AP5m1 protein.

Next, we amplified the wild-type and mutant *ap5m1* cDNA in three overlapping amplicons to sequence the entire coding region (Fig. 4B; Table S1). The middle and 3’ end overlapping PCR products showed no differences in sequence between wild-type and mutant. However, the most 5’ *ap5m1* amplicon amplified successfully from wild type, but was not obtained from mutant cDNA, despite repeated attempts, suggesting a possible splicing defect. Sequencing the 5’ and 3’ ends of intron 1 of *ap5m1* genomic DNA confirmed a splice donor site mutation (G to A) in the exon1-intron 1 junction (Figs. 4C; S1B). This results in predicted retention of the first intron in the transcript. Using primers within the first intron and either exon 1 or exon 2, we found that indeed the *ovy^p37ad^* cDNA retained the intron, resulting in a premature stop codon in the intronic sequence (Figs. 4D and 5A; S1B), expected to cause a loss of all functional domains of the Ap5m1 protein. We then sequenced the *ap5m1* cDNA from the second *ovy^p35aluc^*allele and identified a missense mutation (C to A transversion), changing amino acid threonine 27 to lysine (Figs. 4C and 5A). This residue is conserved in all 9 vertebrate species examined (Fig. 5B), suggesting its importance to Ap5m1 function.

**Figure 5.**
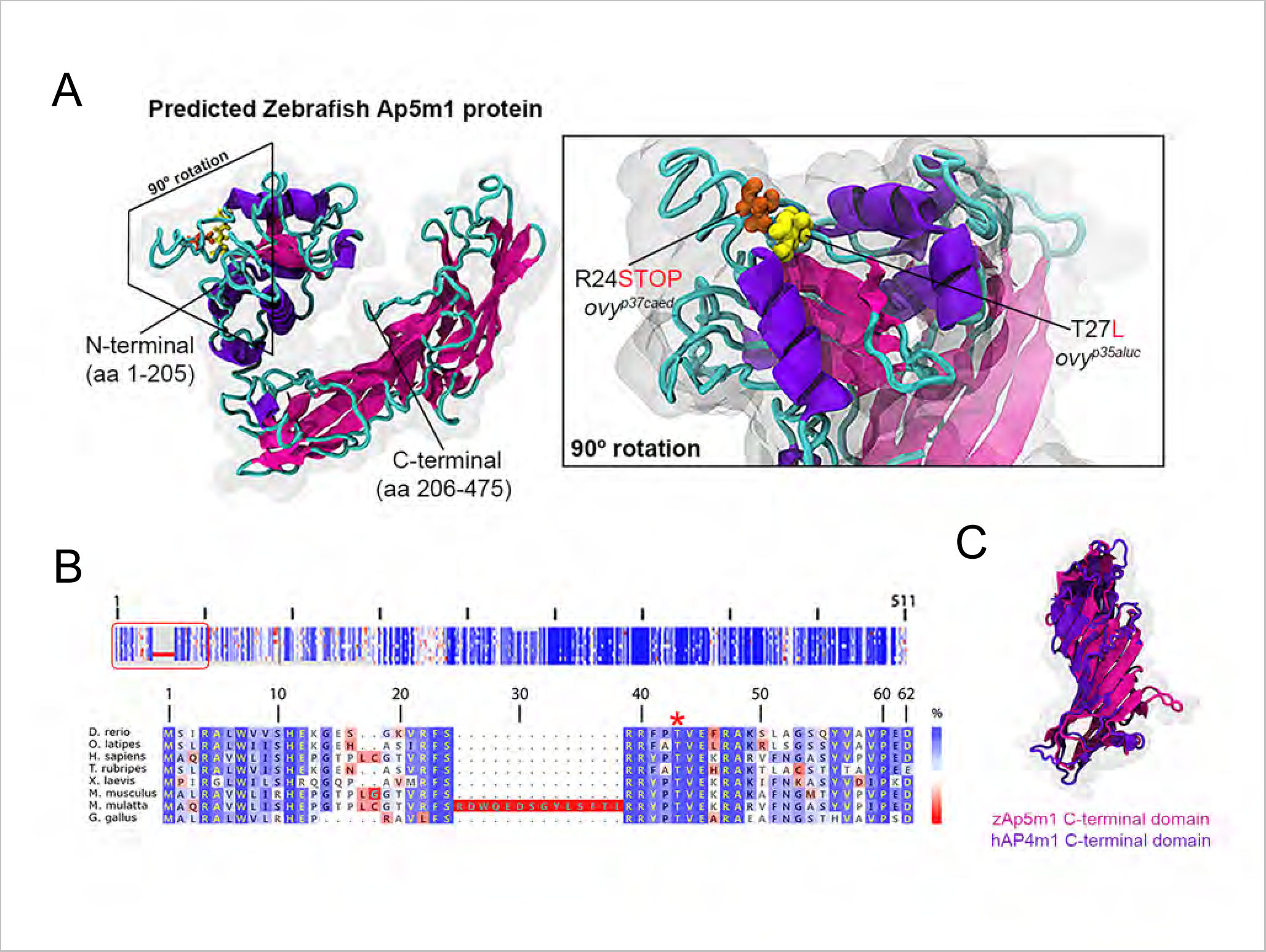
Molecular modeling of the maternal zebrafish AP5m1 factor. **A.** Left: predicted tertiary structure of the zebrafish Ap5m1 protein. The α- and β-helixes are colored in purple and pink, respectively, and the connecting loops in cyan. N- and C- terminal domains are indicated. The boundary of the two domains is at residue 205 (arrow head), which is found in the flexible loop comprised by residues 204-208. The approximate volumetric density map of the protein is shown in transparent gray. Right: predicted domain architecture of the Ap5m1 N-terminal portion containing residues Arginine 24 (R24) and Threonine 27 (T27). The amino acid change and premature stop codon caused by the *ovy^p35aluc^* and *ovy^p37ad^*mutations, respectively, are shown. **G.** Multiple amino acid alignments of the Ap5m1 protein and representative members of the vertebrate lineage. Top: Schematic representation of the protein alignment. The overall percentage (%) identity decreases from top to bottom. The red box indicates the first 49 amino acid residues of Ap5m1. Bottom: Detailed amino acid alignment showing high conservation in fish, amphibians and mammals. Notice high Threonine (T) amino acid conservation (red asterisk), which is mutated in *ovy^p35aluc^*. **H.** Structural superposition between zebrafish Ap5m1 (pink, amino acids 205-475) and the crystal structure of human Ap4m1 C-terminal domains (purple, amino acids 185-453). The RMSD value corresponds to 2.45 Å, thus demonstrating structural identity in overall 3D structure.

### A zebrafish Arp5m1 protein model suggests a function in intracellular trafficking driving yolk globule maturation

To molecularly characterize the Ap5m1 protein, we generated a homology-based structural model of the zebrafish Ap5m1 protein (Fig. 5A). Our model shows that Ap5m1 is comprised of an N-terminal α-helical domain (residues 1-205) and β-sheet structured C-terminal domain (residues 206-492) (Fig. 5A). Immunoprecipitation experiments suggest that the N-terminal domain interacts with another subunit of the AP5 complex (Slabicki et al., 2010). The *ovy^p35aluc^* mutation impacts a peptide loop in the N-terminal domain (Figs. 5A and 5B). The threonine to lysine substitution likely provokes a conformational change that would affect its binding activity. In contrast, the C-terminal or Mu homology domain may be the site of specific cargo recognition and interaction (Hirst et al., 2011). Importantly, the AP5m1 C-terminal domain showed high conservation with acidic residues of the adaptor protein complex 4 mu4 (AP4m4) subunit C-terminal domain and its overall predicted structure (Fig. 5C). AP4m4 is a subunit of the human AP4 complex (another heterotetramer involved in vesicle trafficking), and plays a role in mediating the transport of cargo from the Golgi to endosomes (Burgos et al., 2010; Hirst et al., 2013); thus suggesting that AP5m1 could function in endosomal trafficking during oogenesis.

The morphological and molecular phenotypic features of the *ovy*/*ap5m1* mutant suggest a functional relationship between the endolysosomal route, YG size acquisition, and yolk protein cleavage during zebrafish oogenesis, which was not known previously in egg-laying organisms. Thus, we provide the first phenotypic study in an animal model system of *ap5m1* gene function.

## Discussion

### Molecular genetics of the oocyte-to-egg transition regulation

The mutants reported here and in the companion paper, disrupt genes that are required during oogenesis or activation of the egg, respectively. They genetically uncouple nuclear and cytoplasmic aspects of oocyte maturation, and spatially distributed events of egg activation. In addition, they demonstrate that the process of preparing an egg for embryonic development is not a simple linear series of events. Rather, multiple parallel, coordinated events that do not all depend on successful completion of earlier steps in the process. For example, most of the opaque egg mutants can complete nuclear maturation (meiosis) even though cytoplasmic maturation is compromised.

On the other hand, this same phenotypic class now comprising 7 mutant genes (here and Dosch et al, 2004) all fail in blastodisc formation. Hence, it is possible that this blastodisc defect is a consequence of the failed yolk maturation, however, we cannot exclude that it reflects a secondary function of this class of genes in blastodisc formation. In the former case, two mechanisms could explain the defect in blastodisc formation: i) yolk cytoplasmic maturation is a prerequisite for the reorganization of cytoplasm to the animal pole or ii) the persistence of unmodified or immature yolk interferes with blastodisc formation. Despite an initial successful transition from meiosis to mitosis, the progeny of the mutants reported here do not complete development underscoring the requirement for cytoplasmic maturation to support full developmental potential.

The zebrafish has become a useful model system to identify genetic factors controlling human female reproduction and associated diseases (reviewed in (Hoo, Kumari, Shaikh, Hue, & Goh, 2016). Future genotype-phenotype studies of oocyte and egg biology in this vertebrate model will enhance our knowledge of the maternal gene bio-network regulating the oocyte-to-embryo transition. Importantly, the accessibility to genetic models will illuminate such maternal factors’ biological function in vertebrate reproduction and development, as well as our understanding of the molecular causes of infertility, and eventually improve clinical applications.

### Genetic modulation of YG formation and maturation by egg-laying organisms and its link with human disease

The link between YG size acquisition and cleavage of yolk proteins during embryogenesis has been elegantly investigated in invertebrate animals, including insects, starfish and sea urchin (Finn, 2007; I. Ramos, Machado, Masuda, & Gomes, 2022; I. B. Ramos, Miranda, De Souza, & Machado, 2006; I. B. Ramos et al., 2007; Romano et al., 2004). However, how YGs form and how yolk protein processing is regulated during oogenesis is still unknown in oviparous vertebrates. Here, we identified four maternal factors controlling YG size and yolk protein cleavage during zebrafish oocyte maturation.

The zebrafish maternal-effect mutant genes that cause impaired oocyte cytoplasmic maturation, including cleavage of the major yolk proteins, may have conserved reproductive functions in lower vertebrate species. The small YG phenotype in the *ovy*/*ap5m1* and *p33bjta* mutants suggests that their gene products regulate fundamental events in vesicle trafficking and formation, respectively, during and/or after the yolk protein precursor internalization process of vitellogenesis. In contrast, wild-type-like size of YGs in the *blac^p25bdth^*mutants suggests that these genes might be essential for proper YG protein processing downstream of the *ovy*/*ap5m1* and *p33bjta* function. These studies provide entry points for future investigations of the molecular control of YG biology (Lubzens, Young, Bobe, & Cerda, 2010; Selman, Wallace, & Cerda, 2001). Based on the lysosomal nature of YGs and their proteolytic hydrolase content, it is possible that discovering the molecular regulators of YG formation and function may contribute to our understanding of lysosomal related pathologies in humans (reviewed in (Bonam et al., 2019; Huizing et al., 2008; Platt et al., 2012).

### Functional aspects of egg yolk protein cleavage in animals

Yolk protein cleavage constitutes a critical event in egg-laying species, generating maternal nutrients essential for embryonic growth: the proteolytically cleaved products of Vitellogenin (Vtg), Lipovitellin (Lv) and Phosvitin (Pv) (Wiley & Wallace, 1981). In addition, antimicrobial activity of Pv and its derived cleavage product has been demonstrated *in vitro* and *in vivo,* including evidence in the immunity of zebrafish embryos (Ding, Liu, Bu, Li, & Zhang, 2012; Wang, Wang, Ma, Ding, & Zhang, 2011), which opens new avenues for the study of the immunological role of yolk protein cleavage in oviparous animals during oogenesis.

Although the molecular identification of regulators of yolk biogenesis and proteolytic processing in multicellular organisms has begun to emerge, much of their functional relevance to embryogenesis remains a puzzle (Dosch et al., 2004; Kanagaraj et al., 2014; McNeil & McNeil, 2005; Van Rompay, Borghgraef, Beets, Caers, & Temmerman, 2015); this work). Dissecting the events mediating YG formation and function may help understand: i) conserved or lost functions during oocyte maturation and how different reproductive strategies and nutritional reserves have evolved among vertebrates (Amiel, Leclere, Robert, Chevalier, & Houliston, 2009; Brawand et al., 2008), and ii) the temporal and spatial coordination of oocyte maturation and its link with external sources of oocyte-specific signaling molecules including sexual hormones and growth factors in amniotic and non-amniotic vertebrates.

Advances in female gametogenesis, together with the need to decipher molecular phenotypes during this process, may provide new insights into the mechanisms underlying yolk biology prior to oocyte maturation and the contribution to the study of oviparity-to-viviparity transition in metazoans. In this scenario, knowledge of the underlying molecular mechanisms modulating yolk composition and function in egg-laying organisms might also extend previous findings that link the loss of egg yolk genes and the emergence of new genes in the origin of specific traits such as lactation and placentation (Brawand et al., 2008). Thus, deciphering the molecular circuit controlling yolk biology and function might contribute to understanding the complex cellular energy, nourishment and immunological resource strategies developed in higher organisms during evolution.

### The role of Ap5m1 in reproductive function

The reduced YG size and failure of yolk protein cleavage in the *ovy*/*ap5m1* mutant suggest that Ap5m1 is involved in YG formation and maturation. Thus, Ap5m1 might act in forming large YGs and in regulating in some manner the activity of YG proteases that cleave Vtg during zebrafish oocyte maturation (Lubzens et al., 2010; Selman et al., 2001). Our findings and the Ap5m1 molecular pathway provide evidence of a potential role for Ap5m1 in intracellular trafficking and the sorting processes required for YG biogenesis in oviparous organisms.

At the cellular level, the Ap5 complex acts in intracellular trafficking (Hirst et al., 2013; 2015, 2018). It consists of a heterotetrameric complex including the ζ and β5 (large), μ5 (medium or m5), σ5 (small) subunits (Hirst et al., 2011). Our molecular modeling of the zebrafish Ap5m1 factor and its homology with the human AP4 complex m4 subunit indicate that it may function in endosomal trafficking during zebrafish oogenesis. The *ovy^p35aluc^*T27L missense mutation is expected to disrupt the domain of Ap5m1 that interacts with the Ap5 β subunit, as observed in *in vitro* studies where that same domain has been mutated (Hirst et al., 2013).

The AP5 complex specifically interacts with proteins SPG11 and SPG15/Spastizin in mammalian cell culture. Mutations in these proteins are implicated in Hereditary Spastic Paraplegia (HSP) disease (Hirst et al., 2013; Slabicki et al., 2010). Intriguingly, siRNA-mediated knockdown of Spg11 or Spg15/Spastizin results in the aggregation of early endosomes, consistent with AP5 functioning with these factors in trafficking in mammalian cells (Hirst et al., 2015). Each of the five AP complexes localizes to and functions in distinct subcellular endosomal trafficking compartments, with the Ap5 complex recently shown in cell culture to act at a later point in the endocytic pathway by sorting endosomes back to the Golgi (Hirst et al., 2018). Our *ovy* mutant represents the first multicellular model for *ap5m1* gene function and concomitantly, the Ap5 complex, providing new avenues to study the oogenesis endolysosomal-YG route in a possible vertebrate model of HSP.

### Maternal control of the endolysosomal system during oogenesis

A role for Spg15/Spastizin in reproduction has been reported in zebrafish (Kanagaraj et al., 2014). Maternally deposited Spastizin regulates cortical granule maturation and loss of *spastizin* function causes a similar opaque egg mutant phenotype and defect in yolk protein cleavage as in our *ovy*/*ap5m1* mutant (Dosch et al., 2004; Kanagaraj et al., 2014). Therefore, it will be interesting in future studies to investigate how Ap5m1 and Spastizin function in YG biogenesis and major yolk protein processing.

In this work, we identified a new regulator, the *ap5m1* gene, as a valuable genetic tool for the study of lysosome-related organelles and its function in the endolysosomal pathway in development. Indeed, our results indicate a novel role for Ap5 complex function in female reproduction. Sorting of Cathepsins and ATPases into lysosomes is essential for yolk protein cleavage (Lubzens et al., 2010; Selman et al., 2001). Interestingly, the role of maternal Arl8b protein in Cathepsin-L sorting and lysosomal-mediated degradation of maternal factors in the visceral yolk sac has been described in mammalian embryos (Oka et al., 2017).

Our results highlight the importance of endolysosomal trafficking and protein cleavage during vertebrate embryogenesis. We hypothesize that YG biogenesis relies on Ap5 complex function and its interaction with Spastizin. Epistasis analysis and biochemical approaches involving the sorting of proteases and electrogenic pumps into lysosomes may shed light on Ap5m1-mediated function during oogenesis.

## Acknowledgments

We thank Drs. Leonardo E. Valdivia, Matías Escobar-Aguirre and Matthew Good for critical reading of the manuscript. Dr. Andrea Stout and the CDB (University of Pennsylvania), and CMA Bio-Bio (Universidad de Concepcion) microscopy cores for assistance in the use of the confocal microscopes. Amy Kugath and fish facility staff at the University of Pennsylvania for technical assistance. Andrea Aguilar and Karina Vega-Drake at the Universidad de Concepcion for fish care.

The authors are grateful to the following funding, National Institutes of Health (NIH): R35GM131908, R21HD094096, and R01HD069321 to M.C.M., Becas Chile/Conicyt Proyecto Postdoctorado 74130048, Apoyo FCBI2019-01, Proyecto VRID Investigación Multidisciplinaria 220.031.117-M and ANID Proyecto Fondecyt de Iniciación 11201118 to R.F, Damon Runyon Postdoctoral Fellowship DRG1826-04 to F.L.M., and NIH training grant T32-HD007516 to E.W.A, NRSA postdoctoral fellowships 5F32GM77835 to E.W.A, 1F32GM080926 to L.K; and American Cancer Society postdoctoral fellowships PF-09-125-01-DDC to L.K and PF-05-041-01-DDC to T.G, and NIH grants, R01 HD100035, HD074078, GM103789, and R35 GM122580 to A.G..

## Materials and Methods

### ENU Mutagenesis and fish husbandry

TÜ and AB strain males were mutagenized with 3.3 mM ENU as described in (Mullins, Hammerschmidt, Haffter, & Nusslein-Volhard, 1994). The mutagenesis efficiency was determined by crossing mutagenized males to females that were triple mutant for three mutations that disrupt pigment formation, *albino* (0.04%)*, golden* (0.12%) and *sparse* (0.12%) as in Mullins et al, 1994. Through the natural crossing strategy, we generated 503 F3 families, each bearing two independently mutagenized genomes. The number of genomes screened was calculated according to the formula for pooled F3 embryos, as previously described (Dosch et al., 2004).

Female maternal-effect mutants were identified by crossing F3 females to sibling/cousin or wild-type males and examining their F4 progeny. Phenotyping analysis was performed immediately after egg collection for fertilization and egg activation morphological defects and later at 1-day post fertilization for morphological defects and viability. Females that produced greater than ∼40% phenotypic progeny were retested in crosses with wild-type males to confirm that the phenotype was a strict maternal-effect. No mutants exhibited maternal-zygotic genetics. For further phenotypic characterization and mapping, female maternal-effect mutants that reproducibly produced phenotypic offspring when crossed with a wild-type male were recovered typically from 10-15 intercrosses of the F2 generation of each family. Adult wild-type and mutant fish were maintained for a 14hr light/10hr dark photoperiod at 28 °C.

### Genome mapping, and sequencing for *ovy*/*ap5m1* gene identification

The oocyte-to-egg transition mutations were mapped to a chromosomal locus as described (Dosch et al., 2004; Stickney et al., 2002; Talbot & Schier, 1999). Positional mapping and cloning were carried out through bulk segregant analysis using SSLP markers as described (Knapik et al., 1998). The *ovy^p35aluc^* and *poac^p26ahubb^* mutations were both mapped to Chr 17 but to distinct genetic intervals between 51 and 61 cM, and 58 and 79 cM, respectively (Table 1). By testing 26 females for recombination, the *ovy^p35aluc^*interval was defined to an interval of 35 cM between markers z7170 (2/26 recombinants) and z8980 (7/26 recombinants). The *ap5m1* gene exon 1-intron 1 and intron 1-exon 2 junctions were amplified by PCR using primers targeting the 5’ and 3’ ends of intron 1 (Table S1). The *ap5m1* missense mutation was identified by sequencing the first overlapping gene amplicon from the *p35aluc* mutant background using specific primers (Table S1). The *blac^p25bdth^*and *blac^p26ahubg^* mutant alleles were mapped to the same loci on Chr 5 between 47 and 64 cM (Table 1).

### Gene complementation and PCR-based genotyping

Complementation tests were performed by crossing three *poac^p26ahubb^*heterozygous females to three homozygous *p35aluc* males and two heterozygous *p35aluc* females to two *ovy^p37ad^* homozygous males and raising the progeny to adulthood. Transheterozygous *p35aluc/ovy^p37ad^* females produced 100% mutant opaque egg embryos, whereas transheterozygous *p35aluc/poac^p26ahubb^*females produced all wild-type embryos.

*poac^p26ahubb^*, *blac^p25bdth^* and *blac^p26ahubg^* alleles were genotyped using flanking SSLPs designed for positional cloning. *ovy^p35aluc^*, and *ovy^p37caed^*alleles were genotyped using <u>K</u>Biosciences Competitive <u>A</u>llele-<u>S</u>pecific <u>P</u>CR genotyping system (KASP, KBiosciences) (see Table S1).

### Whole-mount fluorescent *in situ* hybridization and CG staining

Fluorescent *in situ* hybridization of *cycB1* (Kondo, Yanagawa, Yoshida, & Yamashita, 1997) and *dazl* (Maegawa, Yamashita, Yasuda, & Inoue, 2002) was performed in whole-mount oocytes and cryosectioned ovaries (30 µm thick). Oocytes were isolated and fixed according to (Fuentes & Fernandez, 2014).

For CG staining, unactivated and activated eggs were collected from gravid females (Mei, Lee, Marlow, Miller, & Mullins, 2009) and natural crosses, respectively. Eggs were then acid-fixed at 0, 5, 20 and 30 mpa intervals in 5% formaldehyde and 5-8% glacial acetic acid at room temperature, manually dechorionated and placed in gently agitation for 2 hrs, washed in 1X Phosphate-Buffered Saline (PBS) 4 times for 10 min each (Fernandez & Fuentes, 2013). Samples were labeled with fluorescent conjugated *Maclura pomifera* Agglutinin (MPA) diluted to 50 µg/ml as described in (Becker & Hart, 1996, 1999), then washed in 1X PBS 4 times for 10 min each and either placed small petri dishes for *in toto* observation or mounted on glass slides.

### F-actin and meiotic chromosomes labeling

For actin labeling, zygotes were fixed, washed and blocked as in (Becker & Hart, 1999). Then, TRITC-Phalloidin (Sigma) was diluted to 0.5 μg/ml in PBT. Samples were mounted in Vectashield mounting medium with DAPI on glass slides.

### Histology

Ovaries dissected from euthanized females were fixed overnight at 4°C in 4% paraformaldehyde. The next day the fixed tissue was washed in PBS and dehydrated in MeOH. Ovaries were embedded in JB-4 Plus plastic resin (Polysciences) and 5-10 μm sections were cut using a Leica RM 2155 microtome. Sectioned ovaries were stained with Hematoxylin (Sigma-Aldrich), washed in distilled water, stained in Eosin Y (Polysciences), washed in distilled water, and cleared with 50% EtOH. Stained sections were coated with Permount (Fisher), and coverslipped.

### Microscopy, imaging and image processing

Live specimen images were captured using a Leica MZ 12-5 stereomicroscope equipped with a Leica IC80 HD camera and control software on an Apple Macintosh Computer. Histological section images were obtained using a Zeiss AXIOSKOP microscope and ProgRes® Mac CapturePro 2.6 (Jenoptik) software. Fluorescent labeled z-stacks were obtained using a Zeiss LSM 780 or 880 confocal microscope and the Plan-Apochromat 10·/0.45, the Plan-Apochromat 20·/0.8, or the Plan-Apochromat 40·/1.2 immersion water objectives.

Images were processed using Fiji and Adobe Photoshop CC 2017 software.

### SDS-PAGE analysis of major yolk proteins

Ovaries were dissected and sorted oocytes were obtained as described in (Dosch et al., 2004). Sorted oocytes and eggs were homogenized in sodium dodecyl sulfate (SDS) buffer with bromophenol blue and glycerol. Samples were heated at 100 °C for 5 minutes, vortexed, centrifuged, and proteins were resolved on 8% or 12% acrylamide gels and visualized with Coomassie blue.

### ECI, YG diameter, and CG number measurements

To quantitatively examine chorion elevation, the surface area of the egg or chorion was measured. Then, the ratio of the chorion relative to the wild-type 30 mpa egg, which was defined as the egg-to-chorion index (ECI), was determined. The ECI for wild-type eggs was defined as the numerical value for normal chorion elevation (Figs. 1E, 4A).

For YG size quantification, the mean diameters of YGs from wild-type and mutant stage III oocytes were measured on micrographs taken in whole sectioned and stained ovaries (n= 2 wild-type and 2 mutant ovaries). 3 different cytoplasmic regions (DCRs) of 96 x 96 μm (height x width) within the same wild-type (n=12 total DCRs from 4 oocytes) and mutant (n=6 total DCRs per allele from 2 mutant oocytes) oocyte were determined to calculate the length of individual YG diameters.

Wild-type and *krang^p30ahub^* CGs were counted in a determined lateral region (DLR) of 60 x 212 x 212 μm (depth x height x width) in egg z-stacks at 0, 10, 20 and 30 mpa from 2 wild-type and 2 mutant females. For each z-stack, the number of CGs was counted by using the MPA channel and the spot (3 μm diameter) association tool in Imaris. The total number of CGs was calculated by summing up the number of spots associated to single MPA-stained CGs.

Prism 7 (GraphPad) software was used to make graphs and statistical analysis of the data. Samples sizes and statistical values are shown in corresponding figure image or legends. For statistical analysis a Tukey’s multiple comparisons test (Figs. 2C; 3C) was performed.

### Protein sequence analysis and molecular modeling of the zebrafish Ap5m1 and Krang proteins

The similarity of the amino acids in the sequence alignments were scored using the BLOSUM substitution matrix with the software Multiseq (Roberts, Eargle, Wright, & Luthey-Schulten, 2006).

The protein sequences of zebrafish Ap5m1 (Accession XP_005158951.1) and Kiaa0513/Krang (Accession AAH74034, accompanying paper), were searched in public non-redundant databases using iterated PSI-BLAST v.2.2.32+ (3^rd^ iteration), entries with ‘hypothetical’, ‘putative’, ‘modeled’ and ‘environmental samples’ were filtered out using entrez tools (Edgar, 2004). The resulting 270 sequences were filtered with a cut-off minimum of 20% identity and E-score of 1E-5, including full and partial alignments. Most of these sequences consisted of uncharacterized proteins for both Ap5m1 and Krang, respectively. The distance matrix was calculated using MUSCLE v.3.8 (Edgar, 2004).

The Ap5m1 and Krang proteins secondary structure, solvent accessibility, flexibility, among other properties, were determined online using PredictProtein Server (Yachdav et al., 2014). The tertiary structure of Ap5m1 and Krang were modeled online using I-TASSER Protein Structure and Function Prediction Server v.4.3 (Roy, Kucukural, & Zhang, 2010). The Ap5m1 N-and C-terminal domains were defined based on the tridimensional structure of the best model. For Ap5m1 C-terminal domain modeling and structure alignment, the top ranked model based on the crystal structure of AP4m1 (Burgos et al., 2010); Protein Data Bank accession no. 3l81.PDB) was selected. The overall Root Mean Square Deviation (RMSD) of the protein structure alignment between zAp5m1 and hAp4m1 C-terminal domains was 2.45 Å, with an alignment length of 227 amino acids used in the superposition. The structural alignment was made using the TM- align server (Zhang & Skolnick, 2005).

## Supplementary Figure Legends

**Figure S1. Lysis phenotype in the *blac* mutant and intron retention in the *ovy^p37ad^* mutant transcript. A.** Eggs from wild-type, *blac^p25bdth^* and *blac^p26ahubg^* females at 40 mpf. **B.** Sequenced exon1-intron1 and intron1-exon2 junctions of the mutant *ap5m1* cDNA showing the donor splice site mutation, G to A, in intron 1 (black asterisk). mpf, minutes post-fertilization. Scale bar= 1.1 mm (A).

