## Supplemental Table 1 for "Maternal regulation of the vertebrate oocyte-to-egg transition"

**Table S1. Primers for mutants genotyping and *ap5m1* gene mutations sequencing.**

| Primer | Sequence |
|---|---|
| z7170 | Forward Primer: 5'-GGCGAATAGGATTCGACAAA-3'<br>Reverse Primer: 5'-TCATGGAGATGAGTGAGTTGCT-3' |
| z8980 | Forward Primer: 5'-GCCCCAGGGTAACATTTAAC-3'<br>Reverse Primer: 5'-GTTGGGTGCTGGTTTTGACT-3' |
| *ap5m1*.amplicon1 | Forward Primer: 5'-CGGTCAGTCAGTGTTGTTATCG-3'<br>Reverse Primer: 5'-CTACAGCAGAAGGCACCAAT-3' |
| *ap5m1*.amplicon2 | Forward Primer: 5'-AATCCTAGCCTGCCTTCCTCTTG-3'<br>Reverse Primer: 5'- GGTAGGAGCCAAGAATTGGT-3' |
| *ap5m1*.amplicon3 | Forward Primer: 5'-CATTCTTGTCCATCCCTGTGTG-3'<br>Reverse Primer: 5'-GTATCCAGAAGAGACAGGAGC-3' |
| *ap5m1*.5'end | Forward Primer: 5'-CGGTCAGTCAGTGTTGTTATCG-3'<br>Reverse Primer: 5'-GTCTGCTTCTTTCAGGTGAGTG-3' |
| *ap5m1*.3'end | Forward Primer: 5'-TGAATGGCTCTCTGAAGAGG-3'<br>Reverse Primer: 5'-CTACAGCAGAAGGCACCAAT-3' |
| *ovy*$^{p35aluc}$ | 5'-CTCTACATCATCAAAATATTTTTTCTTTGCCTCCTTCAG GCGATTCCCCA[C/A]AGTTGAGTTCCGTGCTAAATCCTTG GCTGGGTCTCATTATGTGGCCGTTC-3' |
| *ovy*$^{p37caed}$ | 5'-TGGGTTGTCTCACACGAAAAGGGTGAATCTGGAAAAG TACGGTTTTCCAG[G/A]TAAGGACCTTGTATATAATTGAC TATTAGCATAGCTGATAGCTTTTGCTA-3' |
