## Supplementary figures and images for "Maternal regulation of the vertebrate oocyte-to-egg transition"

### Supplemental Figure 1

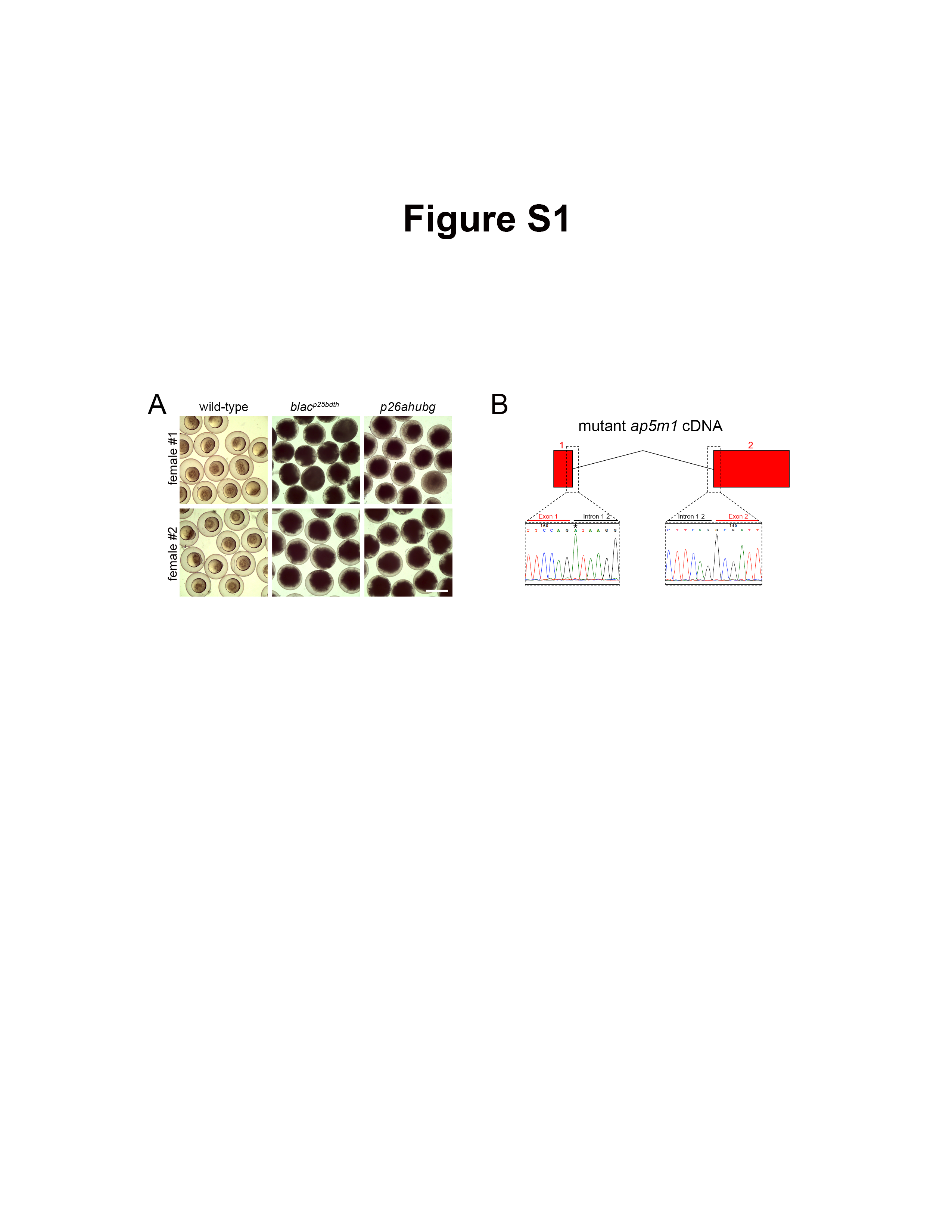
